# The zebrafish dorsolateral habenula is required for updating learned behaviors

**DOI:** 10.1101/802256

**Authors:** Fabrizio Palumbo, Bram Serneels, Robbrecht Pelgrims, Emre Yaksi

## Abstract

Operant conditioning requires multiple cognitive processes, such as learning, prediction of potential outcomes and decision making. It is less clear how interactions of these processes lead to the behavioral adaptations that allow animals to cope with a changing environment. We showed that juvenile zebrafish can perform conditioned place avoidance learning, with an improving performance across development. Ablation of the dorsolateral habenula (dlHb), a brain region involved in associative learning and prediction of outcomes, led to an unexpected improvement in performance and delayed memory extinction. Interestingly, while the control animals exhibited rapid adaptation to a changing learning rule, dlHb ablated animals failed to adapt. Altogether, our results show that the dlHb plays a central role in switching animals’ strategies while integrating new evidence with prior experience.

## INTRODUCTION

The ability to predict threats or rewards, and to shape behavior in response to a changing environment is essential for survival [1]. Operant conditioning has been widely used to study the process of learning, where the likelihood of a behavior is modulated by positive or negative reinforcers. Therefore, operant conditioning requires a constant monitoring of the animal’s own actions, a real time estimation of its consequences and, finally, an update of the relationship between the executed behavior and its outcome [2–4]. The constant update of this relationship between the learned behavior and its outcome is essential, since, in real-life, rules and conditions constantly change. Therefore, a high degree of behavioral flexibility is required in order to maximize survival chances. Reversal learning has been extensively used, across species, to evaluate behavioral flexibility [5–7]. Not surprisingly, the impairment of this behavioral flexibility leads to neuropsychiatric conditions [8] such as obsessive-compulsive behaviors [9], substance abuse [10], post-traumatic stress [11] and mood disorders [9, 12]. Therefore, understanding the neural basis of operant conditioning and the associated behavioral flexibility to learn new rules, is essential to provide solutions to these disorders.

A brain region that plays an important role in operant learning is the habenula (Hb). The lateral habenula (lHb) was shown to encode aversive experiences [13, 14] as well as prediction errors, in case of mismatch between the expectation and the outcome, this both in rodents and primates [15, 16]. Moreover, lesioning the lHb disrupted cost-benefit decisions in rats, without affecting the animal’s ability to evaluate the magnitude of the immediate reward [17]. Complementary studies show the involvement of the medial Hb (mHb) in depression, anxiety, fear and addiction [18–20]. Lesioning the mHb in a rat model of depression, reverses anhedonia-like behavior resulting in a rescued sucrose consumption after chronic stress [21]. Consistent with mammals, several studies highlight the homologous role of the Hb during associative learning in zebrafish. Based on neural connectivity and molecular characteristics, the zebrafish ventral habenula (vHb) and dorsal habenula (dHb) are well-established homologues of the mammalian lHb [22] and mHb [23], respectively. Ablation of the dHb in zebrafish disrupts the execution of experience-dependent fear response [23, 24] and the evaluation of outcomes in social conflict [25]. Whereas, vHb ablation impairs active avoidance learning, without affecting the panic behavior induced by classical fear conditioning [26]. Finally, two recent studies also highlight the different roles of the dHb and vHb in response to recovery after an aversive experience [27] and in the transition of coping behaviors in response to inescapable shock [28], respectively. All these previous studies suggest the involvement of the habenula in coordinating multiple neural processes, evaluating outcomes and selecting optimal behaviors in response to experienced changes in the environment. It is however less clear how changes in learned rules in cognitively demanding tasks are integrated in the brain and what role the habenula plays in this behavioral flexibility. Investigation of such cognitively demanding behaviors in transparent juvenile zebrafish, would open new avenues in studying the neural basis of behavioral flexibility across widely dispersed brain regions.

In this study, we designed an operant conditioning assay to investigate whether transparent juvenile zebrafish can perform conditioned place avoidance (CPA) learning, which requires the coordination of multiple cognitive processes. We showed that while few of the one-week-old larval zebrafish can perform CPA successfully and that animals’ performance improves drastically at the three-week-old juvenile stage with transparent brains. Moreover, we introduced a novel approach to distinguish the operant conditioning from the fear response of the animals. In addition, we showed that the zebrafish CPA performance and memory recall improved drastically across multiple training days. Genetic ablation of the dlHb led to an apparent improvement in performance likely due to delayed memory extinction. Our reversal learning experiments showed that the juvenile zebrafish exhibit prominent behavioral flexibility and can quickly learn changing CPA rules. However, the dlHb ablated zebrafish lacked the behavioral flexibility that is necessary for reversal learning and failed to adapt to changing learning rules. Our results highlight the potential of juvenile zebrafish for performing complex behavioral tasks that require the coordination of multiple cognitive processes and propose a new role for the dlHb in adaptive behaviors.

## RESULTS

### Juvenile zebrafish can perform conditioned place avoidance with increasing performance across development

To investigate the ontogeny of CPA learning, we focus on the first four weeks of development, during which zebrafish are still transparent. We observed that the size of zebrafish increased significantly from larval to juvenile stage (Figure S1B). Using a custom-built behavioral setup (Figure S1A), we trained zebrafish in a CPA protocol, where they received mild aversive electric stimuli only when they entered the conditioned zone of the arena, marked by a red color presented from the bottom. The training was composed of a baseline session followed by two blocks of conditioning and test sessions (Figure 1A, Figure S1A, D). During the baseline, we observed no bias in exploration of the arena (Figure 1A, B, D). During the conditioning sessions, animals received a 10 ms mild electric stimulus every 1.3 seconds, only when they entered the conditioned zone. We quantified learning performance by using multiple measures, such as the reduction of time spent and the distance swum in the conditioned zone, as well as the average distance from the boundary of the conditioned zone. During conditioning sessions C1 and C2, zebrafish exhibited a significant decrease of time spent in the conditioned zone, at all developmental stages (Figure 1A, B, Figure S1F, G). However, only three and four-week-old zebrafish retained a significant avoidance behavior during the entire 30 min long test sessions T1 and T2, with four-week-old animals performing significantly better than three-week-old ones (Figure 1A, B, Figure S1F, G). This result was also confirmed by other measures of learning such as the distance swum in the conditioned zone (Figure 1C, Figure S1F, G) and the animals’ average distance from the conditioned zone (Figure 1D, Figure S1F, G). All groups decreased their swimming velocity during the conditioning protocol (Figure 1E). All groups responded to aversive stimuli with a transient speed increase (Figure S1C). Next, we investigated the animals’ behavior while they approached the conditioned zone. We calculated trajectories before and after reaching the border between the conditioned and the safe zone, during sessions C2 and T2 (Figure 1F, Figure S1E). We observed that several three- to four-week-old animals took evasive action in proximity to the conditioned zone (Figure 1F, gray dots on the right hemisphere, Figure 1G) or never approached the boundary (Figure 1F, black dots). Finally, we showed that the ratio of animals with learning indices significantly higher than chance levels, increased from 31% in one-week-old to above 60% in three- to four-week-old zebrafish (Figure 1H). Altogether, these results show that juvenile zebrafish can successfully perform CPA learning with increasing performance across development.

**Figure 1.**
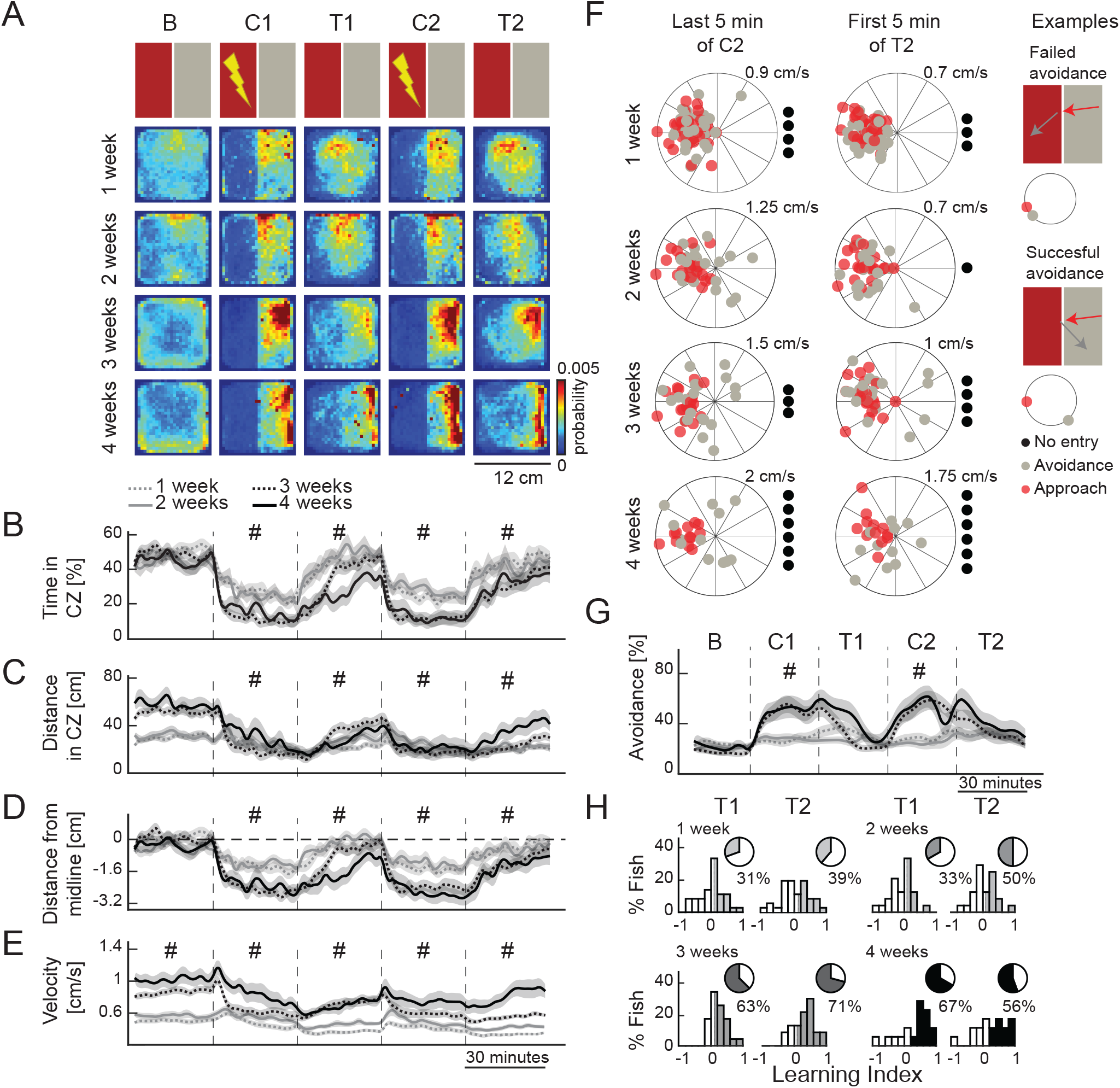
Juvenile zebrafish can perform conditioned place avoidance learning with increased performance across development. (A) The top row shows a schematic representation of the protocol. B: baseline, C1: conditioning 1, T1: test 1, C2: conditioning 2, T2: test 2. Consecutive row shows the heatmaps depicting the average density of zebrafish position for each experimental group (over the course of each entire session), one-week (n=36), two-week (n=24), three-week (n=24) and four-week (n=18) old zebrafish. (B) Percentage of time spent in the conditioned zone (CZ) of the arena. (C) Absolute swimming distance in the CZ of the arena. (D) Average distance of the animals away from the CZ of the arena. Dashed line from 0 indicates the animal being on the midline. (E) Average swim velocity of the zebrafish. (B-E) Line colors and style indicate different age groups. The time course of learning measures in 2 minutes time bins (mean ± SEM). # indicate significance in comparisons across several groups, which are detailed in Figure S1E, F. (F) Zebrafish swimming direction near the midline boundary of CZ and safe zone, during the last 5 min of C2 (left) and first 5 min of T2 (right), increasing age groups are ordered from top to bottom. Each dot on the polar plot depicts the average swim direction of one zebrafish during the four seconds before (red) and after (gray) encountering the midline. For each dot, the distance from the center encodes average swimming velocity of the animal. The numbers on top of the plots indicate the swimming velocity represented by the outer-ring of the polar plots. Black dots outside the polar plots represent animals that never entered the CZ. (G) Average ratio of successful avoidance over the course of the protocol, divided in five-minutes time-bins (mean ± SEM). # indicate significance in comparisons across several groups, which are in Figure S1H. (H) Histograms of the learning indices during the two test sessions, across development. The darker colors for each histogram represent the animals classified as learners, based on the threshold during baseline behavior. The percentage of learners is in the pie chart above each histogram. ***p= <0.001, **p= <0.01, *p= <0.05 Wilcoxon rank sum test. All the statistical comparisons between groups and intra-group are shown in supplementary tables for visualization purpose (Figure S1). Duration of each session is 30min.

### Comparing the fear response with the operant learning behavior

To compare the fear response to aversive stimuli with the operant learning behavior, we designed a sham training protocol for three- to four-week-old, size-matched, juvenile zebrafish. To do this, we trained half of the animals with the CPA protocol, while in parallel a sham group was simultaneously exposed to the same amount of aversive stimuli. To match the aversive experience in both groups, the mild electric stimuli of the sham group were precisely timed and dictated by the actions of their CPA-trained siblings in a neighboring behavioral arena with the same features (color, shape, temperature). Since the sham animals’ aversive experiences were independent of their behavior and solely controlled by the actions of their CPA-trained siblings, the sham group experienced fear in a similar arena without the opportunity to learn (Figure 2A). As expected, the trained zebrafish learned the CPA task and avoided the conditioned zone successfully in all measures of learning (black traces, Figure 2B-D). However, sham animals had no opportunity to learn (red traces, Figure 2B-D). Consequently, we observed that trained animals exhibited a lower average swimming velocity than the sham group (Figure 2E). Hence, the reduction in speed was not merely due to the fear response of the animals, but due to a change in behavioral strategy upon CPA learning. None of the trained or sham animals exhibited substantial freezing behavior (Figure 2F) and both groups responded to the aversive shock with a transient increase in swim velocity (Figure 2G). Moreover, the trained animals strategically took evasive actions along the boundaries between conditioned and safe zones and actively avoided entering the conditioned zone (Figure 2H, I). Sham animals, however, did not perceive these boundaries as relevant features in the environment (Figure 2H, I). These results highlight that juvenile zebrafish exhibit substantially different behaviors during CPA learning, when compared to fear response, and that CPA learning behavior can be separated from fear.

**Figure 2.**
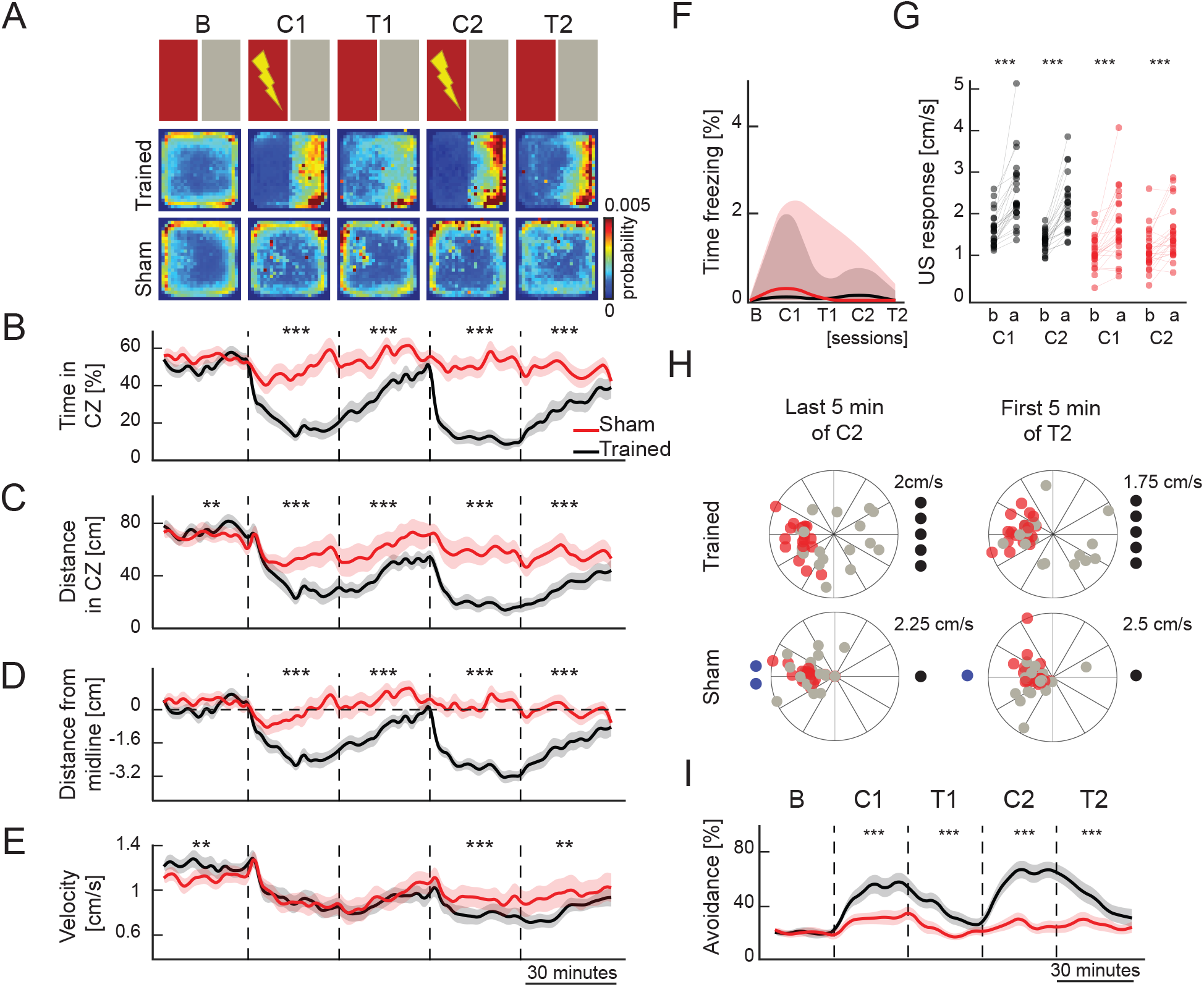
Comparing operant learning with fear response. (A) The top row shows a schematic representation of the protocol. Consecutive rows show the heatmaps depicting the average density of zebrafish position for trained (n=25) and sham (n=25) groups. (B) Percentage of time spent in the conditioned zone (CZ) of the arena. (C) Absolute swimming distance in the CZ of the arena. (D) Average distance of the animals away from the CZ of the arena. Dashed line from 0 indicates the animal being on the midline. (E) Average swimming velocity of the zebrafish. Note that the trained group shows a more prominent reduction of swimming velocity when compared to the sham group. (B-E) Line colors indicate control (black) and sham (red) groups. The time course of learning measures in 2-minute time-bins (mean ± SEM). (F) Percentage of time that the zebrafish exhibit no swimming, which is defined by less then 2mm swimming during 2 seconds. Note the absence of long term freezing for both contol (black) and sham (red) group. Solid lines represent median, shaded areas represent first and third quartiles. (G) Average swimming velocity of the zebrafish, one second before (b) and after (a) the delivery of aversive unconditioned stimuli (US). (H) Zebrafish swimming direction near the midline boundary of CZ and safe zone, during the last 5 min of C2 (left) and first 5 min of T2 (right), in CPA-trained (top) and sham (bottom). Each dot on the polar plot depicts the average swimming direction of one zebrafish during the four seconds before (red) and after (gray) encountering the midline. For each dot, the distance from the center encodes the average swimming velocity of the animal. The numbers on top of the plots indicate the swimming velocity represented by the outer ring of the polar plots. Black dots outside the polar plots represent animals that never entered the CZ, blue dots represent animals that never left the CZ. (I) Average ratio of successful avoidance over the course of the protocol, divided in five-minute time-bins (mean ± SEM). ***p= <0.001, **p= <0.01, *p= <0.05 Wilcoxon signed-rank test. Duration of each session is 30min.

### CPA performance and memory recall improve across multiple training sessions

Next, we investigated whether juvenile zebrafish improve their performance across multiple training days and whether such training facilitates long-term learning and memory consolidation. To achieve this, we trained a set of four-week-old juvenile zebrafish across 4 consecutive days, where each session started with a blank session with no color cues, followed by a recall session with color cues, and two conditioning session interspersed by a test session (Figure 3A). In line with our hypothesis, multi-day training improved the performance in all measurements of learning during the conditioning and test sessions (Figure 3 A-D, F, Figure S2), especially when compared to training day 1 (Figure 3F, Figure S2). In particular, the percentage of animals classified as learners increase from 67%, in day 1, up to 89% in the consecutive training days (Figure 3F). In order to test the memory recall after multi-day training, we analyzed and compared the ten-minute time windows during the switch from blank to the presentation of the recall pattern (boxes labelled B’ and R’ in Figure 3B, C). During this switch, the color pattern of the behavioral arena changes from only grey, to the CPA pattern with red and gray color cues that the animals were trained with (Figure 3A, top). In fact, from the third day of training on, we observed a transient but significant memory recall, when CPA color cues were presented right after the blank session. This was quantified by a reduced time spent (Figure 3G) and a reduced distance swum (Figure 3H) in the conditioned zone. We also observed that during the recall period, the zebrafish perceived the boundary between the color cues and took evasive action as they approached the boundary of the conditioned zone (Figure 3J, K, black arrow). The ratio of successful avoidance during recall significantly increased (Figure 3K) across multi-day training. We did not observe any prominent freezing behavior even after multi-day training (Figure 3I). Our results demonstrated that the CPA performance of zebrafish improves across multiple training days and that zebrafish form long-term CPA memories.

**Figure 3.**
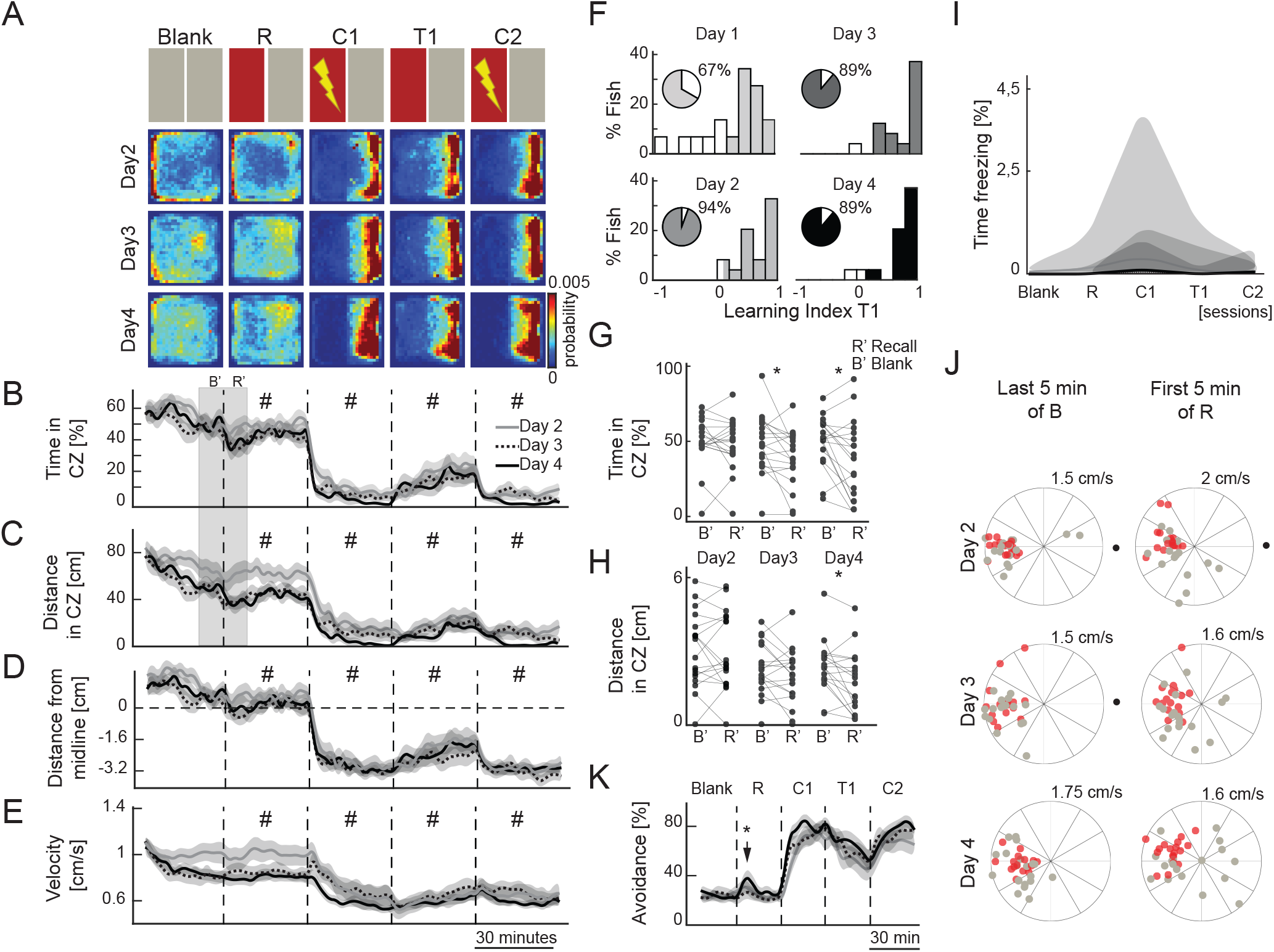
CPA performance and memory recall improve across multi-day training. (A) The top row shows a schematic representation of the protocol. Blank: baseline with no color cues, R: recall, C1: conditioning 1, T1: test 1, C2: conditioning 2. The next rows show the heatmaps depicting the average density of zebrafish position during multiple training days (n=18). (B) Percentage of time spent in the conditioned zone (CZ) of the arena. (C) Absolute swimming distance in the CZ of the arena. (D) Average distance of the animals away from the CZ of the arena. Dashed line from 0 indicates the animal being on the midline. (E) Average swimming velocity of the zebrafish. (B-E) Line colors and style indicate zebrafish performance during multiple training days. The time course of learning measures in two-minute time-bins (mean ± SEM). # indicate significance in comparisons across several groups, which are detailed in Figure S2A. Note the improved performance over consecutive training days. (F) Histograms of the learning indices during the test sessions, across multiple days of training. The darker colors for each histogram represent the animals classified as learners, based on the threshold during baseline behavior. The percentage of learners is in the pie chart above each histogram. (G) Average time spent by the animal in the CZ during the last ten minutes of the blank session (B’) and in the first ten minutes of the recall session (R’) over multiple training days. B’ and R’ periods are marked with the grey box in panel B and C. Note that from the third training day on, zebrafish exhibit aversion to CZ when the color cues are presented during recall ression. (H) Average swimming distance in the CZ during the last ten minutes of the blank session (B’) and in the first ten minutes of the recall session (R’) over multiple training days. B’ and R’ periods are marked with the grey box in panel B and C. Note that on the fourth day of training zebrafish exhibit aversion to CZ when the color cues are presented during recall ression. (I) Percentage of time that the zebrafish exhibit no swimming, which is defined by less then 2mm swimming during 2 seconds. Note the absence of long-term freezing for any of the training days. Solid lines represent median, shaded areas represent first and third quartiles. (J) Zebrafish swimming direction near the midline boundary of CZ and safe zone, during the last ten minutes of Blank (B’, left) and first ten minutes of Recall (R’, right), consecutive training days are ordered from top to bottom. Each dot on the polar plot depicts the average swim direction of one zebrafish during the four seconds before (red) and after (gray) encountering the midline. For each dot, the distance from the center encodes the average swimming velocity of the animal. The numbers on top of the plots indicate the swimming velocity represented by the outer ring of the polar plots. Black dots outside the polar plots represent animals that never entered the CZ. (K) Average ratio of successful avoidance over the course of the protocol, divided in five-minute time-bins (mean ± SEM). # indicate significance in comparisons across several groups, which are detailed in Figure S2. Note the improved performance over consecutive training days. ***p= <0.001, **p= <0.01, *p= <0.05 (B, C, D, E) Wilcoxon rank-sum test, (G, H, K) Wilcoxon signed-rank test. All the statistical comparison between groups and intra-group are shown in supplementary tables for visualization purpose. Duration of each session is 30min.

### Dorsolateral habenula ablation leads to an apparent improvement of CPA performance

Dorsolateral habenula ablation in adult zebrafish was shown to interfere with the execution of experience-dependent fear responses [23]. We hypothesize that dlHb ablation might also interfere with CPA learning. We achieved chemogenetic ablation of the dlHb by incubating three to four-week-old juvenile zebrafish Tg(narp:GAL4VP16; UAS-E1b:NTR-mCherry) with 10mM metronidazole (MTZ) (Figure 4A, B). In juvenile zebrafish, narp:Gal4 is expressed exclusively in dlHb neurons, and the expression pattern is comparable to the one in adult zebrafish [23] (Figure S3 A-F). The slightly more lateral location of narp:Gal4 expressing dlHb neurons in adults is due to sequential habenular neurogenesis across development [29]. Next, we compared dlHb ablated juvenile zebrafish with the control group in a multi-day training protocol. To our surprise, we observed that dlHb ablated animals outperformed controls already at the end of the first day of training, and that their CPA performance remained higher than the control group over the consecutive training days (Figure 4C-N). From the third day of training on, we also observed that dlHb ablated animals exhibited robust CPA recall behavior (Figure 4O, P). Importantly, we did not observe prominent freezing behavior in dlHb ablated animals even after several days of CPA training (Figure S3G). These results suggest that dlHb ablation leads to an apparent improvement of CPA performance and a better memory recall, across multi-day training sessions.

**Figure 4.**
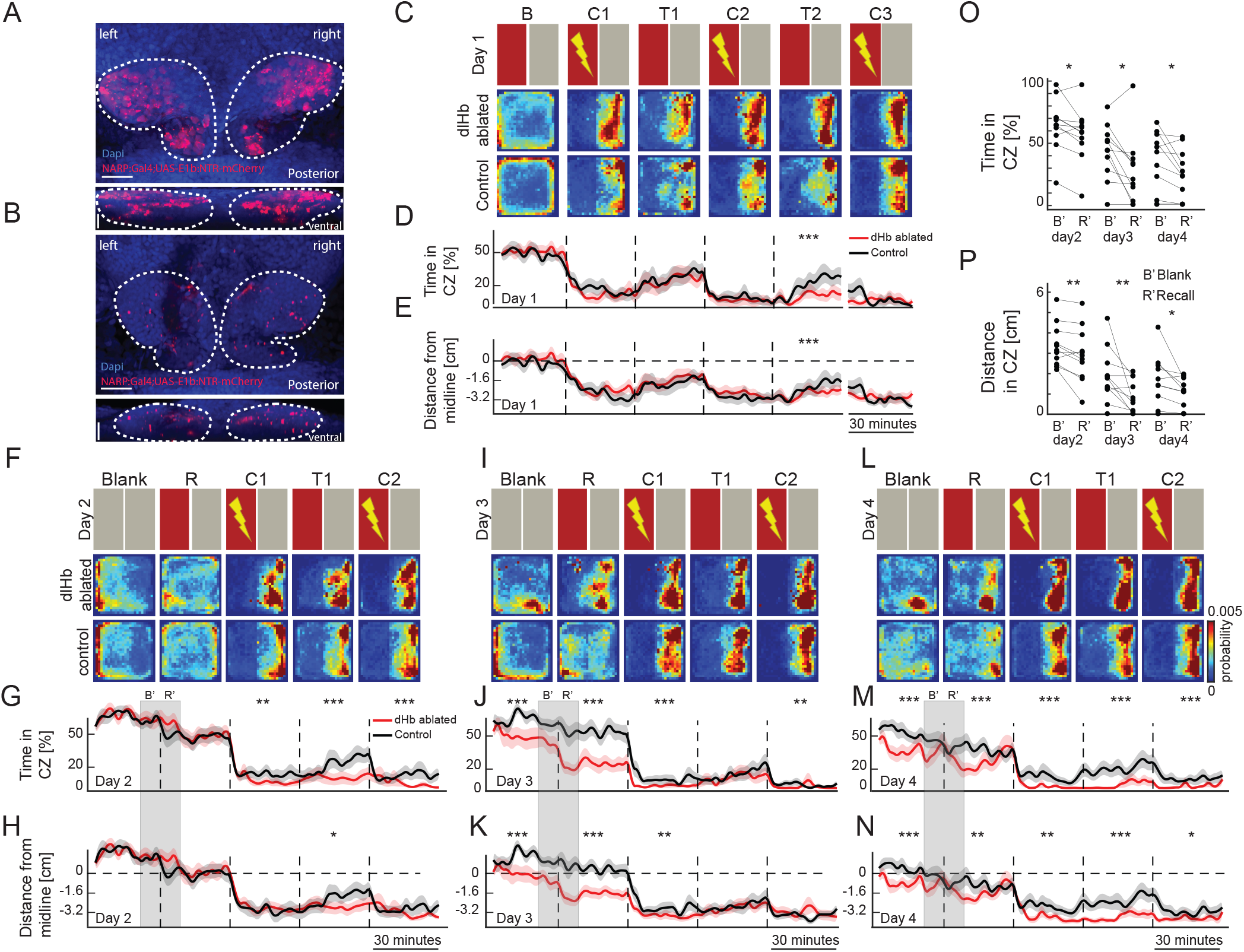
Dorsolateral habenula ablation improves CPA performance and delays memory extinction. (A,B) Confocal microscopy images of Tg(narp:Gal4;UAS-E1b:NTR-mCherry) zebrafsh before (A) and after (B) Metrodiniazole (MTZ) treatment. Scale bar is 25 μm. (C,F,I,L) The top row shows a schematic representation of the protocol. Blank: baseline with no color cues, R: recall, C1: conditioning 1, T1: test 1, C2: conditioning 2. The next rows show the heatmaps depicting the average density of zebrafish position for dlHb-ablated (n=11) and control (n=12) groups, across multiple training days. (C) First day of training. (F) Second day of training. (I) Third day of training. (L) Fourth day of training. (D, G, J, M) Percentage of time spent in the conditioned zone (CZ) of the arena, during multiple training days for dlHb-ablated (red) and control (black) zebrafish. Note the improved performance for dlHb-ablated zebrafish during the recall and test sessions. (E, H, K, N) Average distance of the animals from the CZ of the arena, during multiple training days for dlHb-ablated (red) and control (black) zebrafish. Dashed line from 0 indicates the animal being on the midline. Note the improved performance for dlHb-ablated zebrafish during the recall and test sessions. (O) Average time spent by the dlHb-ablated zebrafish in the CZ during the last ten minutes of the blank session (B’) and in the first ten minutes of the recall session (R’) over multiple training days. Note that already from the second training day on zebrafish exhibit aversion to the CZ when the color cues are presented during the recall session. (P) Average swimming distance of the dlHb-ablated zebrafish in the CZ during the last ten minutes of the blank session (B’) and in the first ten minutes of the recall session (R’) over multiple training days. Note that already from the second day of training zebrafish exhibit aversion to the CZ, when the color cues are presented during the recall session. ***p= <0.001, **p= <0.01, *p= <0.05 (D, E, G, H, J, K, M, N) Wilcoxon rank-sum test, (O, P) Wilcoxon signed-rank test. Duration of each session is 30min.

### The dorsolateral habenula is important for behavioral flexibility during reversal learning

We hypothesized that one reason for dlHb ablated zebrafish to perform better than control animals can be an impairment in memory extinction. To test this hypothesis, we introduced an extended extinction session (T3) after the previously described CPA training. In line with our hypothesis, the dlHb ablated zebrafish showed a significantly slower extinction rate of the avoidance behavior, for the entire duration of the extended extinction session (Figure 5A-C). The slower memory extinction in dlHb animals can in principle be explained by an inefficient integration of new information with past experience. Since the dlHb ablated animals performed better during the initial CPA protocol (Figure 4 and 5), we hypothesize that they did not integrate new information about the lack of aversive experience. Inspired by these results, we next reversed the learning rule after the initial CPA training, and paired the aversive stimuli with the part of the CPA arena marked with a grey visual cue (Figure 5, conditioning periods C3 and C4). In this reversal learning protocol, zebrafish were required to update their learned CPA behavior, and relearn to associate the initially safe zone with the aversive experience. Excitingly, the dlHb ablated zebrafish did significantly worse than control animals in reversal learning, quantified by all indicators of reversal learning performance (Figure 5A-D). We observed no significant differences in the velocity of dlHb ablated versus control group (Figure 5E) as well as no prominent freezing behavior (Figure S4A). Both groups significantly increased their transient swimming velocity upon administration of the aversive stimulus (Figure S4B). Altogether our results reveal that while dlHb ablation leads to a better performance in a CPA protocol, dlHb ablated zebrafish exhibit impaired ability to integrate new evidence into their past experience when tested in a memory extinction and reversal learning paradigm.

**Figure 5.**
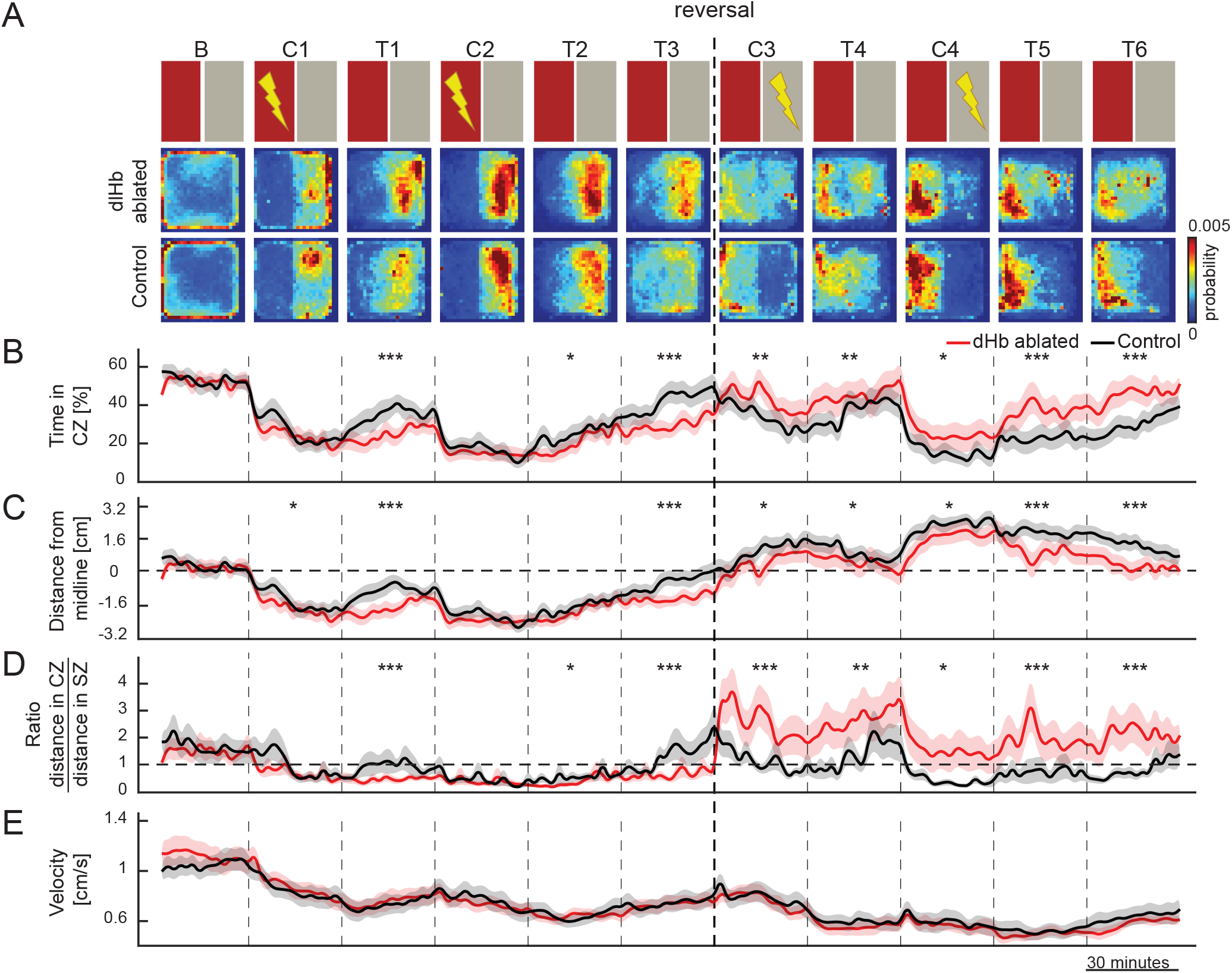
The dorsolateral habenula is important for behavioral flexibility during reversal learning. (A)The top row shows a schematic representation of the protocol. B: baseline with color cues, C1: conditioning 1, T1: test 1, C2: conditioning 2, T2: test 2, T3: test 3 (extinction test), C3: conditioning 3 (reversal learning), T4: test 4 (reversal learning test), C4: conditioning 4 (reversal learning), T5: test 5(reversal learning test), T6: test 6(reversal learning extinction test). The next rows show the heatmaps depicting the average density of zebrafish position for control (n=21) and dlHb-ablated (n=22) groups. (B) Percentage of time spent in the conditioned zone (CZ) of the arena. (C) Average distance of the animals away from the CZ of the arena. Dashed line from 0 indicates the animal being on the midline. (D) The ratio of swimming distance in conditioned zone (CZ) versus safe zone (SZ). (E) Average swimming velocity of the zebrafish. (B-E) Line colors indicate control (black) and dlHb-ablated (red) groups. The time course of learning measures in two-minute time-bins (mean ± SEM). ***p= <0.001, **p= <0.01, *p= <0.05, no mark=non-significant, Wilcoxon rank-sum test. Duration of each session is 30min.

## DISCUSSION

Series of recent studies highlight the increasing popularity of zebrafish for studying diverse aspects of animal behavior [30–39], including complex behaviors such as learning [25, 26, 40–46] and social interactions [47, 48]. While studying complex behaviors in adult zebrafish allows investigation of brain function in a fully mature brain, larval and juvenile zebrafish have the tremendous advantage of a small and transparent brain, when it comes to studying brain activity [33, 38, 49–58]. Hence, we investigated the ontogeny of a complex operant behavior, namely CPA learning, and showed that the learning performance of young zebrafish improves across development. Our findings are in line with a previous study [45], but also show that even at younger ages, some zebrafish can perform such CPA learning. At the third week of development the learning performance of juvenile zebrafish reaches very high levels and becomes more robust across animals. Interestingly, we observed that the CPA learning strategy of younger (one- to two-week-old) animals are substantially different than older (three- to four-week-old) animals. While young animals swim away from the aversive stimuli towards the safe zone in the conditioning sessions, during the test sessions the visual boundary between the conditioned zone and the safe zone has no impact on their behavior. Older animals, however, perceive the visual boundary and adopt swimming strategies to take evasive action, in order to avoid the conditioned zone during the test session. Moreover, after multi-day training, four-week-old juvenile zebrafish can recall those memories even after 24hr. Adult zebrafish were shown to recall memories encoded by telencephalic neural activity [43], homologous to the mammalian cortex [59], resembling the memory engrams described in mammals [60–62]. Our results showed, for the first time, such long-term memory recall in juvenile zebrafish with small and transparent brains that are amenable for brain-wide optical imaging, at single neuron resolution. Future studies will be necessary to further investigate the neural basis of memory at different time scales, and to understand how brain regions that are involved in short-term and long-term learning interact with each other to generate memory engrams.

The major brain structures required in operant learning, such as the amygdala and the hippocampus, are evolutionarily conserved across vertebrates [44, 59, 63–68]. However, all these brain regions mature during vertebrate development by the addition of new cell types, new layers or even entire brain structures [69–71]. For instance, most neurons in the zebrafish homolog of the amygdala [44] are born later at the juvenile stage but not during the larval stage [65]. Similarly, mammalian entorhinal-hippocampal circuitry matures gradually as animals develop [69, 72]. A recent study in the zebrafish telencephalon revealed a sequential addition of distinct neural layers across development [73], which likely represents the developmentally expanding functionality of this ancestral cortico-limbic circuitry. Our findings can therefore reflect an overall maturation of the zebrafish forebrain allowing animals to integrate new types of information about their environment with their experiences and evaluate them in a context-specific framework to take critical decisions.

An increasing number of studies use fear learning for studying the molecular and neural basis of learning [60–62, 74–76]. However, it is not always easy to distinguish the role of fear from learning, in modifying molecular and neuronal processes in the brain. Here, we introduced a novel approach, for comparing and perhaps distinguishing fear responses from learning. We propose that one way to address this challenge is to simultaneously train control animals in a CPA protocol while comparing the results to a sham group. This sham group experiences the same environment and the exact same aversive stimuli at the same time as the control group, but without a CPA rule to learn. We argue that comparing the control group to the sham group, allow us to separate the learning-related processes in the brain from all other processes related to the subjective experiences of the animals. For example, our data suggest that the sham animals did not develop a preference for a particular zone of the arena or its visual features. We also observed that the swim patterns or speed of the sham animals were not modulated as prominently as in those that were allowed to learn. Finally, we also observed no freezing behaviors, neither in trained, nor sham animals. This suggests that our CPA protocol induces only a very mild aversive experience. Likely due to a very low aversive stimulus intensity, we observed no helplessness behaviors or freezing [23, 27, 28] even after several days of CPA training. Altogether, we propose that using very mild aversive stimuli and comparing CPA trained animals with sham animals is a promising approach to be used in future studies for disentangling the neural processes underlying learning from fear, anxiety, and helplessness.

Our results suggest that CPA-trained animals pay attention to the visual boundary between the conditioning zone and the safe zone. They use this salient information and they take evasive action once they approach it, in order to avoid the conditioned zone. In our multi-day training experiments, we also observed that zebrafish can store such visual information for several days and avoid the conditioned zone, as soon as the color cues are presented. These results suggest that CPA learning in juvenile zebrafish is a complex behavior that relies on the integration of salient information with emotional responses to short- and long-term aversive experiences, while making appropriate decisions, similar to mammals[77, 78]. Previous studies in mammals established a direct role of the habenula in the prediction of learned outcomes [13, 15, 16, 26, 79, 80], and negative motivational values [14, 81, 82]. Similarly, the zebrafish dorsal and ventral habenula were implicated in classical and operant learning [23, 24, 26]. At first glance, our results for improved CPA performance in dlHb ablated zebrafish were surprising. In fact, dlHb ablated zebrafish also outperformed the control animals in memory recall session during the early period of multi-day training. After several days of training, the control animals caught up with dlHb ablated animals in CPA performance and memory recall, highlighting the amount of training as an important factor while interpreting these results. It is also important to note that due to the low intensity of the aversive stimuli, neither control nor dlHb ablated animals showed any significant freezing or helpless behavior [23, 24], which might have explained the apparent improvement of CPA performance. Instead, we observed that the dlHb ablated animals showed a significantly slower memory extinction rate. In fact, the extinction of a memory is an essential component of learning new rules [83]. The extinction of fear memories was proposed as a therapeutic target for PTSD treatment [84]. Hence, comparing predicted outcomes with the actual consequences is important for updating the previous rules and memories with the new ones [5, 85]. Our reversal learning experiments show that juvenile zebrafish exhibit such behavioral flexibility and can quickly adapt to changing rules. However, we also showed that the dlHb ablated zebrafish with slower memory extinction have difficulty adapting to changing CPA rules during reversal learning. In line with our findings, knockout mice lacking the narp gene that is expressed both in the mouse mHb and the zebrafish dlHb, were shown to succeed in instrumental conditioning but failed in a devaluation protocol, which required adjustment of choices in the absence of rewards[86]. Primate studies highlights further roles for the habenula in adaptive behaviors [16, 85]. In human subjects, impaired behavioral flexibility can lead to various mood disorders [87–90]. In line with all these studies, our findings highlight an important role for the habenula in optimizing animal behavior to provide the necessary behavioral flexibility, when there is a mismatch between the predicted and the actual outcomes. In the future, it will be crucial to investigate the neural computations across widely dispersed brain regions of transparent juvenile zebrafish, that could underlie the role of the habenula and its input/output structures in behavioral flexibility and learning.

## ACKNOWLEDGEMENTS

We thank H. Okamoto (RIKEN Center for Brain Science, Japan), M. Ahrens (HHMI, Janelia Farm, USA) for transgenic zebrafish lines. We thank S. Eggen, M. Andresen, V. Nguyen, A Nygard and our fish facility support team for technical assistance. We thank Nathalie Jurisch-Yaksi and Stephanie Fore for helpful comments on the text. We thank all Yaksi lab members for stimulating discussions. This work was funded by ERC starting grant 335561 (F.P., R.P., E.Y.) and RCN FRIPRO Research Grant 239973 (E.Y.). Work in the E.Y. lab is funded by the Kavli Institute for Systems Neuroscience at NTNU.

## AUTHOR CONTRIBUTIONS

Conceptualization, F.P., E.Y.; Methodology and data, F.P., B.S.; Recording software F.P., R.P.; Data Analysis, F.P.; Investigation, all authors; Writing, F.P., E.Y.; Review & Editing, all authors; Funding Acquisition and Supervision, E.Y.

## METHODS

### Experimental animals

The animal facilities and maintenance of the zebrafish, Danio rerio, were approved by the NFSA (Norwegian Food Safety Authority). Fish were kept in 3,5 liter tanks in a Techniplast Zebtech Multilinking system at constant conditions: 28°C, pH 7 and 600μSiemens, at a 14:10 hour light/dark cycle to simulate optimal natural breeding conditions. Fish received a normal diet of dry food (SDS 100-400, dependent of age) two times/ day and Artemia nauplii once a day (Grade0, platinum Label, Argent Laboratories, Redmond, USA). For the experiments investigating the ontogeny of CPA learning, the age range in each group had a variability of plus or minus two days. For all the experiment shown in Figure 1, 2 and 3 we used Tg(elavl3:GCaMP6s) [91] zebrafish. In the experiment shown in Figure 4 and 5 we used Tg(narp:GAL4VP16; UAS-E1b:NTR-mCherry)[23] and siblings not expressing NTR-mCherry protein as controls. All experimental procedures performed were in accordance with the directive 2010/63/EU of the European Parliament and the Council of the European Union and the Norwegian Food Safety Authorities.

### Behavioral setup

Custom-made real-time tracking software was implemented to track six zebrafish simultaneously, using the OpenCV 3.0 library and the QtCreator5.2 developing platform. A microcontroller controls six integrated circuits to deliver electrical stimulation to each individual arena. Each current pulse has a duration of 10 ms and an amplitude of approximately 1.2 mA. This results in a current density of approximately 0.1 mA/cm using tungsten electrodes. A pulse of current is delivered every 740 ms, resulting in a delivery frequency of 1.33 Hz. The aversive stimulus is administered to the fish only when it is located inside the conditioned zone. The visual stimulus is presented from the bottom of the arena using a horizontally positioned LCD monitor. Each of the six arenas was divided into 2 equal parts with red and grey colors with matched luminosity. A Manta 235B camera was used to monitor animal behavior at 15 fps. Experiments were conducted in Gosselin square Petri dishes 120mm × 120mm × 15.8 mm with opaque side walls. The entire behavioral setup was enclosed in a temperature-controlled black isolation box to prevent any external interference. The water temperature during training was kept at 26°C. A schematic of the behavioral setup is shown in Figure S1A.

### Experimental procedures

One fish per arena was placed in the behavioral setup between 09:00 and 11:00 am, after the morning feeding, and the experiment was started immediately. Every animal was used only in one specific protocol once. All the behavioral protocols performed in this work are composed of the following sessions that were used in various combinations:

Baseline session: the animal’s position was monitored and recorded for 60 minutes, while the color pattern was displayed to the animal.

Blank session: position was monitored and recorded for 30 minutes, but no color pattern was presented to the animal; the arena was entirely covered by a uniform gray color.

Conditioning session: position was monitored and recorded for 30 minutes. Only when the zebrafish enters the conditioned zone a mildly aversive weak (10 ms, 1,2 mA) electric stimulus was delivered at 1,33 Hz.

Test session: position was monitored and recorded for 30 minutes, while the color pattern was displayed to the animal.

In Figure 1, 2, 3 and 4, the “conditioned zone” was always associated with the left half of the arena. In Figure 1,3 and 4 the conditioned side of the arena was always associated with the red color. In the experiment presented in Figure 2 half of the animals were trained against the red color pattern, the other half against the gray color pattern. We observed no substantial differences in this two groups and combined both groups for data analysis. Each of these conditions had the corresponding sham control animals. The reversal protocol in Figure 5 was achieved by first conditioning the animals against the left/red side of the arena. During the third and fourth conditioning session the aversive stimulus was experienced only on the right/gray side of the arena.

### Metronidazole treatment

To ablate the neurons of dlHb, Tg(narp:GAL4VP16; UAS-E1b:NTR-mCherry) zebrafish were treated with 10mM metronidazole (MTZ, Sigma-Aldrich) for 24h followed by a washout period of at least 12h, protocol adapted from Agetsuma et al. (2010) [23]. The treatment was started two days prior the behavioral protocol. The fish were placed, in groups of three, in a Petri dish containing 50ml of 10mM MTZ +250 microliters of DMSO. Control group was subjected to the same treatment. The fish were kept in the 28°C incubator, covered in aluminum foil, for 24h. The next day the fish were transferred to a new Petri dish containing fresh artificial fish water and they were fed with dry food and placed back in the incubator. The water was changed at least one additional time before the experiment.

### Behavioral analysis

To perform all the behavioral analysis shown in this study, we used custom-made scripts written in MATLAB. The heat-maps are an average of zebrafish swimming position over the entire length of each session. For all further analysis, of the baseline session only the last 30 minutes are taken into consideration. We divided each behavioral session into 2-minute time bins to be able to better detect changes over time.

The learning index is defined as follows:

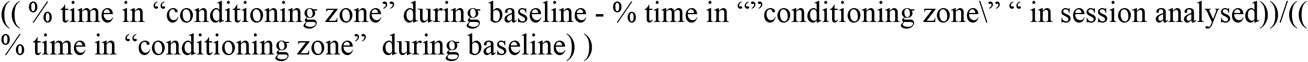

To discriminate between learners and non-learners, a lowest boundary for the learning index was calculated, by using the relatively random distribution of zebrafish swimming position during the baseline session. All fish that exceeded this lowest boundary were considered to exhibit significant learning upon CPA training. Forquantifying swimming direction and avoidance behaviors, the position was detected when fish were near the midline boundary between conditioned zone and safe zone. Next, the linearized trajectory of the animal in the four seconds preceding and following the encounter with the midline boundary were used as approach and avoidance, respectively. We then quantify, in 5-minute time bins, the successful avoidance rate of the animals. If the position of the animals was in the “conditioned zone” after they encountered the midline the avoidance was considered unsuccessful, otherwise it was considered successful.

In reversal learning experiments, we only continued the reversal protocol with those animals that exhibited prominent CPA learning in the first training session and classified as learners using our learning index criterion. If animals did not learn the initial CPA task, they were not subjected to reversal learning.

Choice of statistical analysis:

The Wilcoxon rank sum test was used to quantify statistical differences between two different populations of fish. The data points in all time-bins were pooled together within each session for each experimental group. The statistical test was then performed between the obtained distributions and compared across experimental groups. When comparing intra-group measurement over the course of the protocol, the entire baseline period was used in order to have a robust estimate of baseline behavior on a population level. Then, each time bin was compared to this baseline distribution.

The Wilcoxon signed-rank test was performed when individual behavior was compared between two different conditions.

### SUPPLEMENTARY MATERIALS

**Figure S1.**
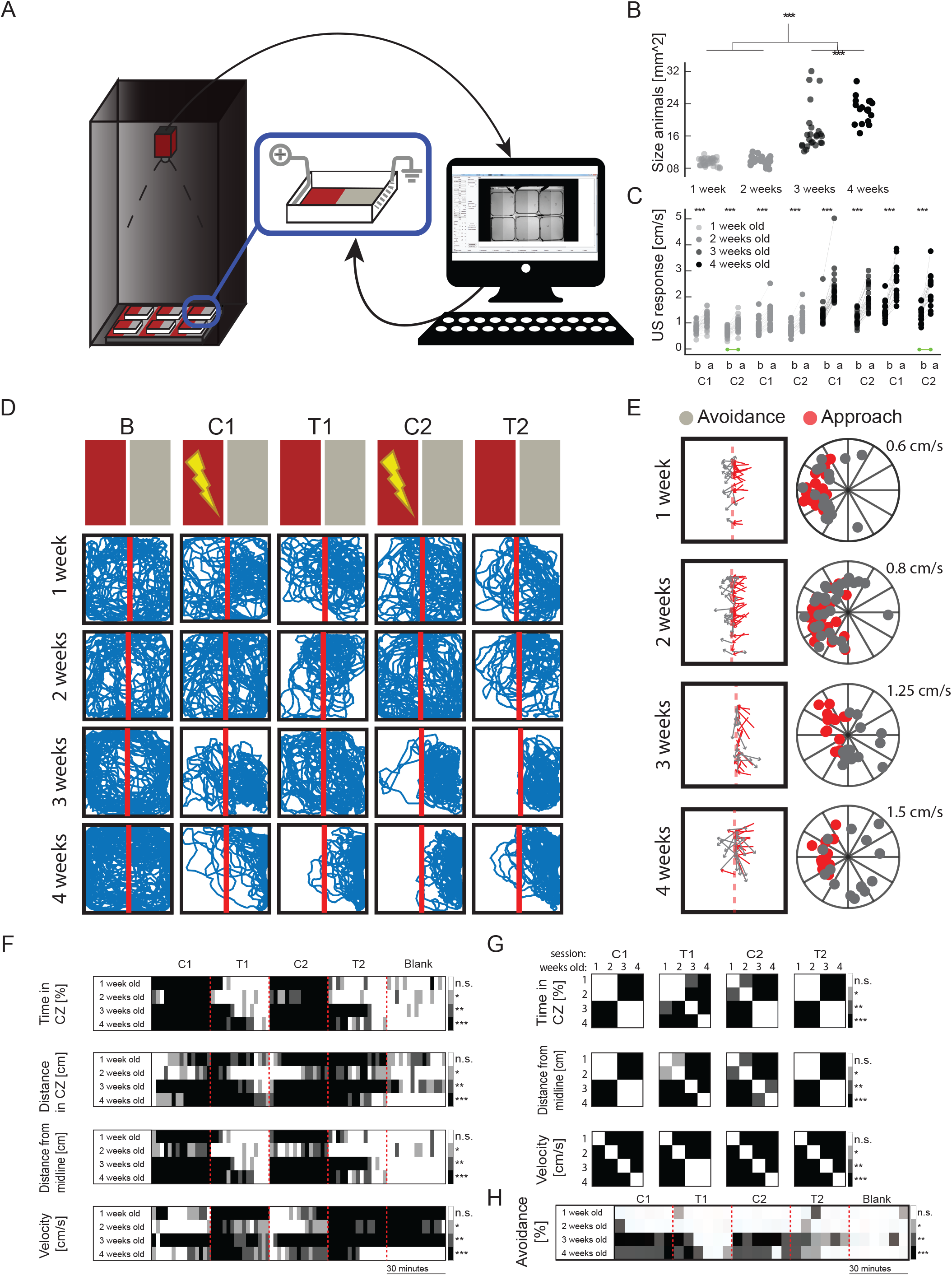
Conditioned place avoidance (CPA) setup and additional analysis of physiological and behavioral parameters during training and multi-group comparisons. (A) Graphical representation of the CPA training setup. (B) Quantification of animal size reveals a significant enlargement of developing zebrafish at 3-4 weeks. ***p= <0.001, Wilcoxon rank-sum test. (C) Average swimming velocity of the zebrafish, one second before (b) and after (a) the delivery of aversive unconditioned stimuli (US). Green dots represent animals never receiving an US. ***p= <0. 001, Wilcoxon signed-rank test. (D) Individual example, for all age groups, showing raw traces of single animal behavior over the course of the protocol. (E) Individual example, for all age groups, showing raw traces of single animal behavior for every single approach of the conditioned zone for the entire duration of the second test session. (left column) Raw traces of approaches (red) and avoidances (gray) of the conditioned zone. (right column) Polar plots representing the example shown by the left column of the panel. (F) Tables showing intra-group statistical comparison of the parameters measured of animal behavior dividing each session in two-minute time-bins. ***p= <0.001, **p= <0.01, *p= <0.05, Wilcoxon rank-sum test. (G) Tables showing statistical comparison across the 4 experimental groups. ***p= <0.0005, **p= <0.005, *p= <0.05, Wilcoxon rank-sum test. (H) Tables showing intra-group statistical comparison of the successful avoidance performed by the animal. ***p= <0.001, **p= <0.01, *p= <0.05, Wilcoxon rank-sum test.

**Figure S2.**
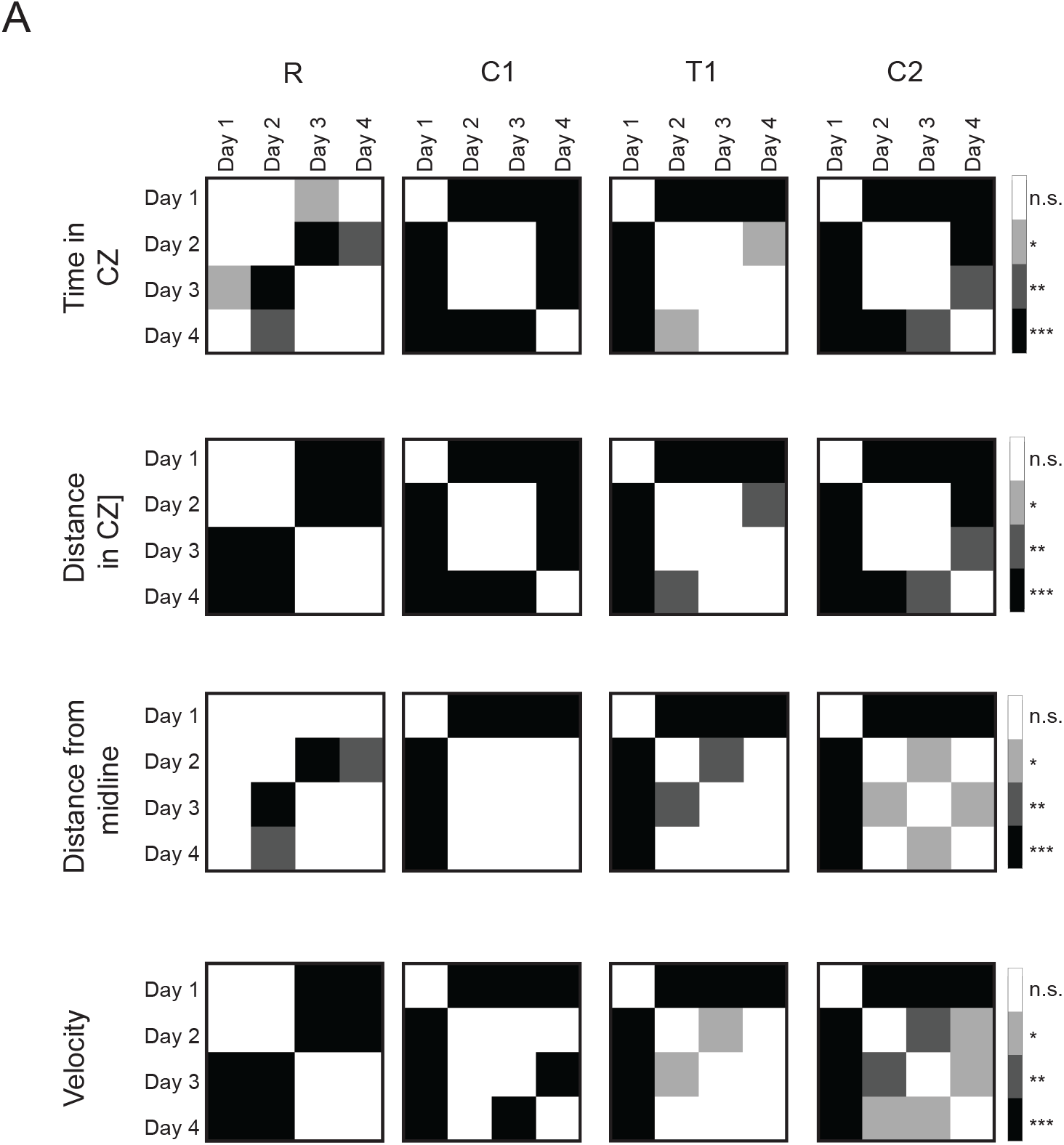
Multi-day training significantly improves performance of the animal. (A) Statistical comparison across the different days of training. R: recall, C1: conditioning 1, T1: test 1, C2: conditioning 2. Note the significant improvement of learning performance (in T1) and recall (R) over consecutive training days, using different learning measures. White color marks non-significant comparison. ***p= <0.001, **p= <0.01, *p= <0.05, Wilcoxon rank-sum test.

**Figure S3.**
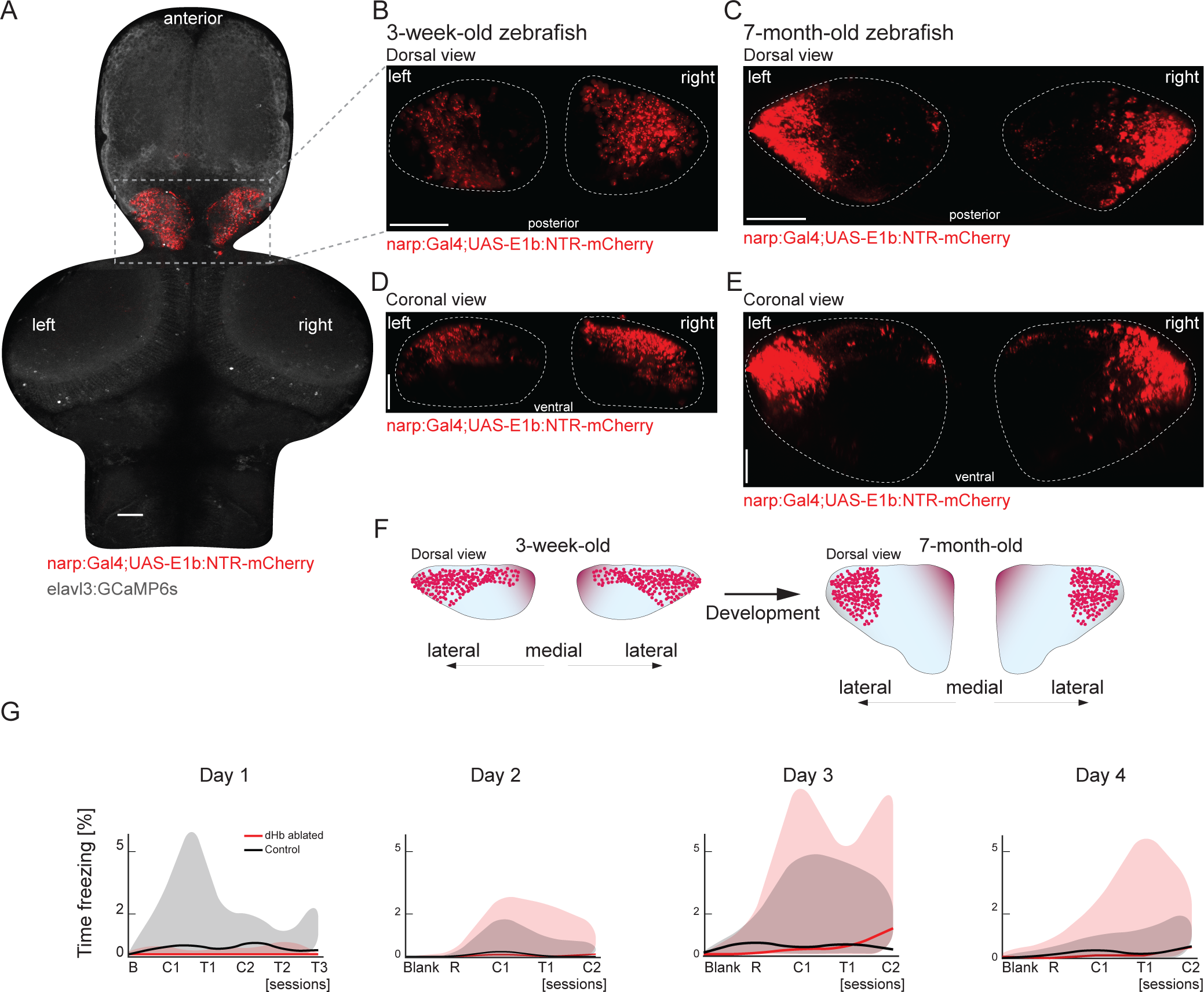
Dorsolateral habenula (dlHb) ablation does not lead to freezing behavior over multi-day training. (A)Confocal microscopy image of the entire brain in three-week-old Tg(narp:Gal4;UAS-E1b:NTR-mCherry;elavl3:GCaMP6s) zebrafish, dorsal view, scale bar is 50μm. Please note that the narp:Gal4 expression is exclusively in the dorsolateral habenula. (B-E) Confocal microscopy images of the habenula in three-week- (B, D) and seven-month-old (C, E) Tg(narp:Gal4;UAS-E1b:NTR-mCherry) zebrafish. Dorsal views (B, C) and coronal views (D, E). Scale bars are 50μm. (F) Schematic representation of narp:Gal4-expressing dorsal habenular neurons in juvenile and adult zebrafish, coronal view. Please note that the relative position of dorsal habenular neurons labelled with narp:Gal4 expression moves towards the lateral end of dorsal habenula, in line with sequential neurogenesis of habenular neurons across development, which was described by Fore et al 2019 (G) Percentage of time that the zebrafish exhibit no swimming, which is defined by less than 2mm swimming during 2 seconds. Note the absence of long-term freezing for both control (black) and dlHb-ablated (red) group. B: baseline, Blank: baseline with no color cues, R: recall, C1: conditioning 1, T1: test 1, C2: conditioning 2, T2: test 2, T3: test 3. Solid lines represent median, shaded areas represent first and third quartiles. dlHb-ablated animals are displayed in red, control in black.

**Figure S4.**
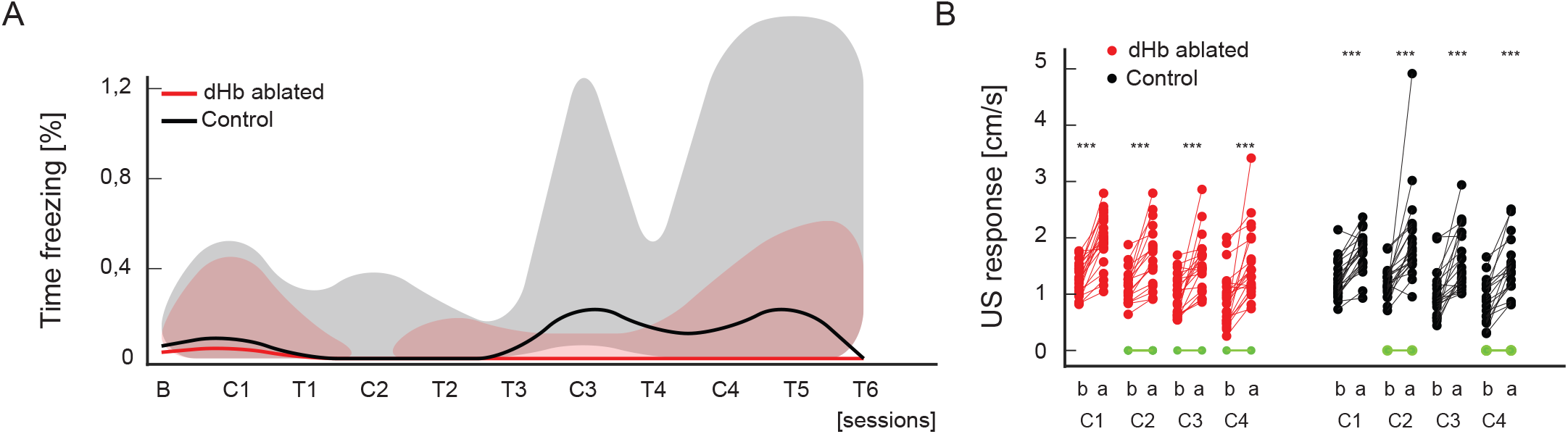
Dorsolateral habenula (dlHb) ablation does not elicit freezing behavior and does not affects animal responses to the unconditioned stimulus during the CPA reversal learning protocol. (A) Percentage of time that the zebrafish exhibit no swimming, which is defined by less than 2mm swimming for 2 seconds. Note the absence of long-term freezing for both control (black) and dlHb-ablated (red) group. B: baseline with color cues, C1: conditioning 1, T1: test 1, C2: conditioning 2, T2: test 2, T3: test 3 (or extinction), C3: conditioning 3 (reversal learning), T4: test 4 (reversal learning test), C4: conditioning 4, T5: test 5, T6: test 6. Solid lines represent median, shaded areas represent first and third quartiles. (B) Average swimming velocity of the zebrafish, one second before (b) and after (a) the delivery of aversive unconditioned stimuli (US). Green dots represent animals never receiving an US. ***p= <0.001, **p= <0.01, *p= <0.05, Wilcoxon signed-rank test. dlHb-ablated animals are displayed in red, control in black.

